# Maternal gut *Bifidobacterium breve* modifies fetal brain metabolism in germ-free mice

**DOI:** 10.1101/2023.12.31.573756

**Authors:** Jorge Lopez-Tello, Raymond Kiu, Zoe Schofield, Douwe van Sinderen, Gwénaëlle Le Gall, Lindsay J Hall, Amanda N Sferruzzi-Perri

## Abstract

In recent years, our understanding of the gut microbiome’s impact on host physiology and metabolism has grown exponentially. Yet, the specific role of certain microorganisms in regulating gestational health and fetal development remains largely unexplored. During pregnancy, *Bifidobacterium* represents a key beneficial microbiota genus that provides multiple benefits, including changes in placental development and fetal glycaemia. In this study, using germ-free mice colonized with or without *Bifidobacterium breve* UCC2003 during pregnancy, we demonstrated that this bacterium is important for controlling fetal brain metabolism. In particular, presence of maternal *Bifidobacterium* led to reduced levels of ten metabolites (including citrate, 3-hydroxyisobutyrate, and carnitine) in the fetal brain, with concurrent elevated abundance of transporters involved in glucose and branched-chain amino acid uptake. *B. breve* supplementation was also associated with increased expression of critical metabolic and cellular pathways, including the PI3K-AKT, AMPK, STAT5 and Wnt-β-catenin (including its receptor Frizzled-7) in the fetal brain. Furthermore, maternal-associated *Bifidobacterium* resulted in HIF-2 protein stabilization and altered a number of https://pubmed.ncbi.nlm.nih.gov/38269505/ genes and proteins involved in cellular growth, axogenesis, and mitochondrial function. These findings highlight that *Bifidobacterium breve* colonisation of the maternal gut is important for the metabolism and growth of the fetal brain.

## Introduction

Fetal growth restriction (FGR) is a severe condition defined as the failure of the fetus to reach its growth potential due to pathological compromise. A main cause of FGR is placental insufficiency during gestation (Malhotra et al., 2019). Human epidemiological studies and experimental animal models have found that FGR and placental insufficiency can affect fetal organ development, including changes in key organs like the heart, kidneys and brain (Brown and Hay, 2016; Camm et al., 2018; Gilchrist et al., 2021; Hinchliffe et al., 1991; López-Tello et al., 2017a; López-Tello et al., 2017b; Luna et al., 2016; Saha et al., 2014). Studies have also shown that disruption of fetal brain development due to suboptimal intrauterine environments can lead to neurodevelopmental disorders postnatally, including motor and cognitive dysfunctions, learning impairments and cerebral palsy (Benítez-Marín et al., 2021; Guellec et al., 2011; Mikaelsson et al., 2013; Miller et al., 2016; Morsing et al., 2011; Vossbeck et al., 2001). Managing placental insufficiency and FGR in a clinical setting can present substantial challenges. Pharmacological interventions like aspirin, heparin and sildenafil citrate, which target inflammatory, coagulation and blood flow pathways are controversial due to variability in their effects on feto-placental and pregnancy outcomes (Bettiol et al., 2019; Pels et al., 2020). Hence, there is an imminent need to devise effective treatments that can prevent and/or mitigate the adverse consequences associated with FGR.

In recent years, there has been an explosion of studies describing the importance of the gut microbiota in regulating developmental processes, from neurogenesis (He et al., 2020) to ageing (Bárcena et al., 2019; O’Toole and Jeffery, 2015). Moreover, a perturbed gut microbiota has been linked to neurological conditions, like Parkinson’s disease (Sampson et al., 2016) and schizophrenia (Zhu et al., 2020), and metabolic-related disorders, including type-2 diabetes (Gurung et al., 2020). In the context of pregnancy, previous studies have demonstrated dramatic changes in the composition of the maternal gut microbiota during pregnancy (Koren et al., 2012) and in women who developed a hypertensive disorder of pregnancy, preeclampsia or had abnormal placental growth (Huang et al., 2021; Miao et al., 2021). Of note, the genus *Bifidobacterium* increases in abundance in the maternal gut during pregnancy in both women and mice (Nuriel-Ohayon et al., 2019) and has been shown to possess multiple benefits. For example, *Bifidobacterium* protects against infectious diseases and is involved in modulating host immune responses (Hidalgo-Cantabrana et al., 2017; Hughes et al., 2017; Kiu et al., 2020a; Kiu et al., 2020b; O’Callaghan and van Sinderen, 2016). Moreover, we have previously shown that administration of *Bifidobacterium* (specifically, *B. breve* UCC2003) to pregnant mice improves fetal growth together with beneficial structural and functional alterations in the placenta and changes in metabolic genes in the fetal liver (Lopez-Tello et al., 2022). Driven by these previously obtained insights, herein we tested the hypothesis that maternal gut *B. breve* abundance would associate with changes in the development and metabolism of the fetal brain. Working with germ-free mice that lack a maternal microbiota (GF) and GF mice that received *B. breve* UCC2003 treatment during pregnancy, our results demonstrate the significance of maternal gut *B. breve* in fetal brain development and metabolism.

## Results

### Maternal gut B. breve UCC2003 modifies expression of genes involved in cell cycle and axogenesis

In our previous publication, we reported that three doses of *B. breve* UCC2003 on gestational days 10,12 and 14, to pregnant GF mice (group referred throughout the text as BIF; Figure S1) improved fetal weight and liver size compared to untreated GF mice. However, brain weight did not differ between untreated GF and BIF fetuses, suggesting that although untreated GF fetuses were growth restricted, they exhibited preserved brain development (Lopez-Tello et al., 2022). To understand the molecular basis of this, we quantified the mRNA levels of key growth, cell cycle, microglia, and neurogenesis genes in the brain of fetuses from GF mice following maternal *B. breve* UCC2003 administration. This revealed that there was no difference in the expression of vascular gene *Vegf* or apoptotic genes *Tp53*, *Casp3* and *Bax* in the brain between fetuses of the GF and BIF groups (Figure 1A). However, mRNA levels of the transcriptional activator *Foxm1* and the mitotic cycling *Cdk1* gene were significantly reduced in the fetal brain of the BIF compared to the untreated GF group (Figure 1B). No changes were observed in two other assessed mitotic cell cycle genes (*Cdk2* and *Cdk4*) or in genes involved in microglia activation (e.g. *Ier3*, *Klf2* or *Egr1)* between GF and BIF groups (Figure 1B-C). Quantification of expression levels of key axonogenesis genes revealed that *Plxna3* was significantly down-regulated in the fetal brain of BIF compared to untreated GF mice (Figure 1D). Also, *Slit1* showed a tendency to be reduced in the BIF treated group (p=0.05). The expression levels of three other axonogenesis genes assessed (*Sema3f*, *Ntn1* and *Nrcam*) were shown to be similar between the two experimental groups (Figure 1D). Collectively, these data demonstrate that maternal gut *B. breve* UCC2003 modulates the expression of genes involved in cell cycle control and axonogenesis in the fetal brain.

**Figure 1.**
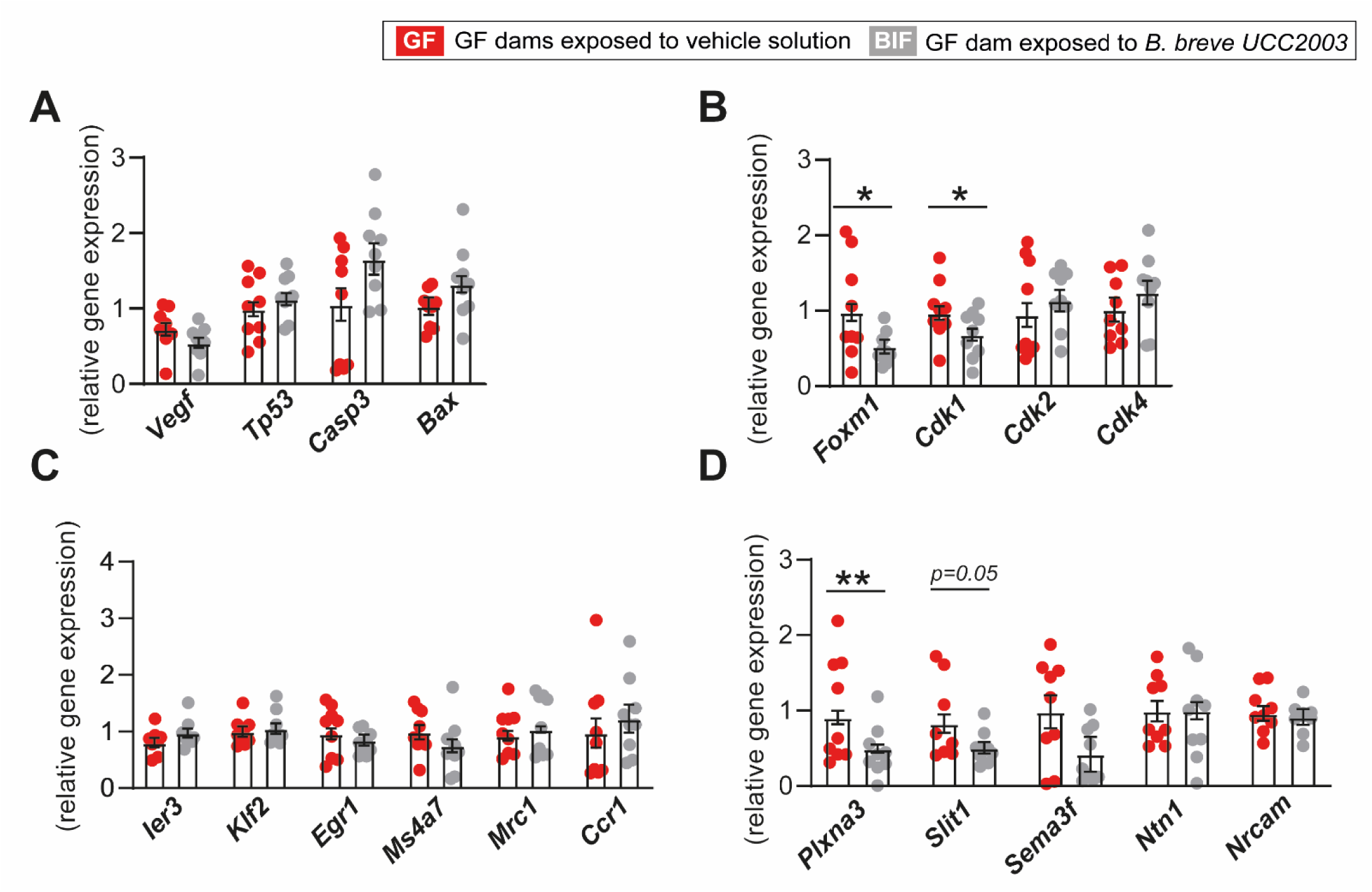
Maternal gut *B. breve* supplementation modifies genes involved in fetal brain cell growth and axon development. Relative mRNA levels of genes involved in cell cycle (A-B), microglia activation (C), and axonogenesis (D). Gene expression is relative to two housekeeping genes (*Gapdh* and *Actb*). Data analysed by ANOVA with the group as fixed effect and means comparisons was made by Fisher test (further details in the methods section). Data are means±SEM with individual data points shown. \**P* < 0.05; \*\**P* < 0.01.

### Maternal gut B. breve UCC2003 modifies fetal brain metabolism

To investigate if maternal *B. breve* UCC2003 supplementation affects fetal brain metabolism, we performed metabolomic profiling of fetal brain lysates from untreated and BIF treated GF mice. We quantified a total of 78 metabolites, of which 10, primarily amino acids and citrate were significantly reduced in the BIF compared to untreated GF group (Table 1 and Table S1). Considering that the levels of certain amino acids such as leucine and valine were altered in the BIF group, we quantified the mRNA levels of transporters involved in branched-chain amino acid uptake and metabolism in the fetal brain. This analysis identified that expression of the large neutral amino acid transporter 1 (LAT1), encoded by the *Slc7a5* gene, but not large neutral amino acid transporter 2 (LAT2; encoded by *Slc7a8*) was significantly elevated in the BIF treated group compared to untreated GF mice (Figure 2A). No differences were found in the expression levels of genes encoding other amino acid transporters, namely the system A amino acid transporters (*Slc38a1,2,4 - SNATs*) in the fetal brain between BIF versus untreated GF mice (Figure 2B).

**Figure 2.**
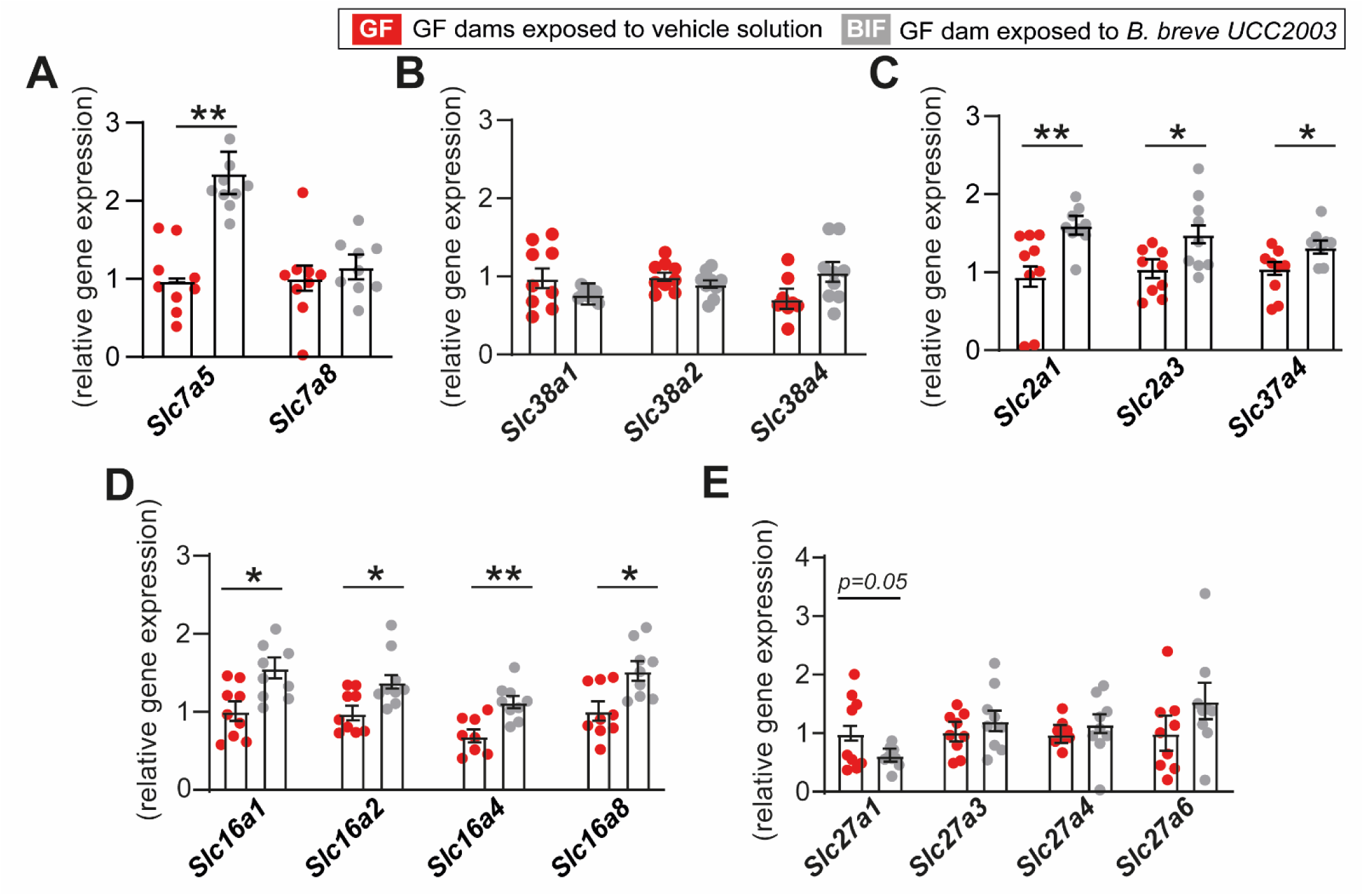
Maternal gut *B. breve* supplementation modifies the expression of nutrient transporters in the fetal brain. (A-E) Relative mRNA levels of genes encoding nutrient transporters measured in fetal brain samples, expression is relative to two housekeeping genes (*Gapdh* and *Actb*). Statistical analysis performed by ANOVA with the group as fixed effect (each fetus as a repeated measure) and means comparisons was made by Fisher test (linear mixed model). Data are means±SEM with individual datapoints shown. Data obtained from a total of 5 GF and 6 BIF pregnant dams/litters (9 fetuses per group analysed). \**P* < 0.05; \*\**P* < 0.01.

**Table 1.** Maternal gut *B. breve* supplementation induce changes in fetal brain metabolites. Data analysed by one-way ANOVA, with the group as fixed effect and means comparisons made by Fisher test (general linear model-GLM model). Litter size added as a covariate. Data displayed as mean±SEM. Values were considered statistically significant with P<0.05. Further data about the metabolites analysed can be found in Table S1.

| Metabolite (mmol/Kg) | GF group (n=5) | BIF group (n=5) | P value |
| --- | --- | --- | --- |
| 3-Hydroxyisobutyrate | 0.063 ± 0.004 | 0.041±0.007 | 0.048 |
| Alanine | 3.523±0.578 | 1.776±0.163 | 0.020 |
| Arginine | 0.192±0.030 | 0.107±0.015 | 0.045 |
| Aspartate | 1.937±0.228 | 1.119±0.150 | 0.026 |
| Carnitine | 0.370±0.047 | 0.210±0.023 | 0.022 |
| Dimethylamine | 0.481±0.192 | 0.072±0.053 | 0.002 |
| Leucine | 0.261±0.019 | 0.177±0.017 | 0.027 |
| Threonine | 2.209±0.292 | 1.170±0.249 | 0.020 |
| Valine | 0.378±0.048 | 0.262±0.027 | 0.035 |
| Citrate | 0.219±0.025 | 0.154±0.023 | 0.046 |

In previous work we reported that circulating levels of glucose were increased in response to *B. breve* UCC2003 administration of GF pregnant mice (Lopez-Tello et al., 2022). Glucose is the predominant source of energy used by the fetal brain to support its development (Vannucci and Vannucci, 2000). Hence, we quantified the mRNA levels of the main glucose transporter-encoding genes *Slc2a1* and *Slc2a3* (GLUT1 and GLUT3, respectively) in fetal brain tissue. This revealed that expression of both glucose transporters was significantly elevated in the fetal brain of BIF treated mice compared to untreated GF group (Figure 2C). Moreover, the mRNA level of *Slc37a4* (encoding G6PT), an important gene that controls glucose homeostasis by regulating the transport of glucose-6-phosphate from the cytoplasm to the lumen of the endoplasmic reticulum (Pan et al., 2009), was also elevated in the BIF group (Figure 2C). In addition, the expression of solute carrier family 16 members 1,2,4,8 - MCTs which are involved in the movement of monocarboxylates, including lactate and pyruvate, and ketone bodies like D-β-hydroxybutyrate (Pérez-Escuredo et al., 2016), were all shown to be significantly elevated in the fetal brain of BIF treated pregnant mice compared to the untreated GF group (Figure 2D). There was no difference in the expression of genes involved in lipid uptake (solute carrier family 27 members - FATP) in brain lysates of BIF treated pregnant GF mice compared to untreated GF mice (Figure 2E). These findings indicate that the presence of maternal gut *B. breve* UCC2003 impacts on the levels of specific metabolites and expression of key nutrient transporters in the fetal brain.

### Maternal gut B. breve UCC2003 modifies cellular and metabolic pathways in the fetal brain

Driven by the changes detected in metabolites and nutrient transporters, we proceeded to analyze cellular and metabolic pathways in the fetal brain. Between the BIF and untreated GF groups, at gene level, we observed elevated levels of *Mapk1* (encoding mitogen-activated protein kinase 1), reduced *Stat5b* (signal transducer and activator of transcription 5B), and unchanged expression of *Pkb* and *Prkaa1* (encoding AKT serine/threonine kinase 1 and protein kinase AMP-activated catalytic subunit alpha 1, respectively) in the fetal brain (Figure 3A). Immunoblotting analysis of PI3K-AKT, MAPK, AMPK and STAT5 signalling pathways (Figure 3B) revealed increased levels of PI3K-p110β and AMPK proteins in fetal brain lysates of BIF treated compared to untreated pregnant GF mice (Figure 3C). Informed by phosphorylation levels, we observed enhanced activation of the AKT signalling pathway (indicated by increased threonine 308 residue phosphorylation) and STAT5 pathway (with increased tyrosine 694 residue phosphorylation) (Figure 3D). Collectively, these data show that the presence of maternal gut *B. breve* UCC2003 is associated with the activation of key cellular and metabolic pathways in the fetal brain.

**Figure 3.**
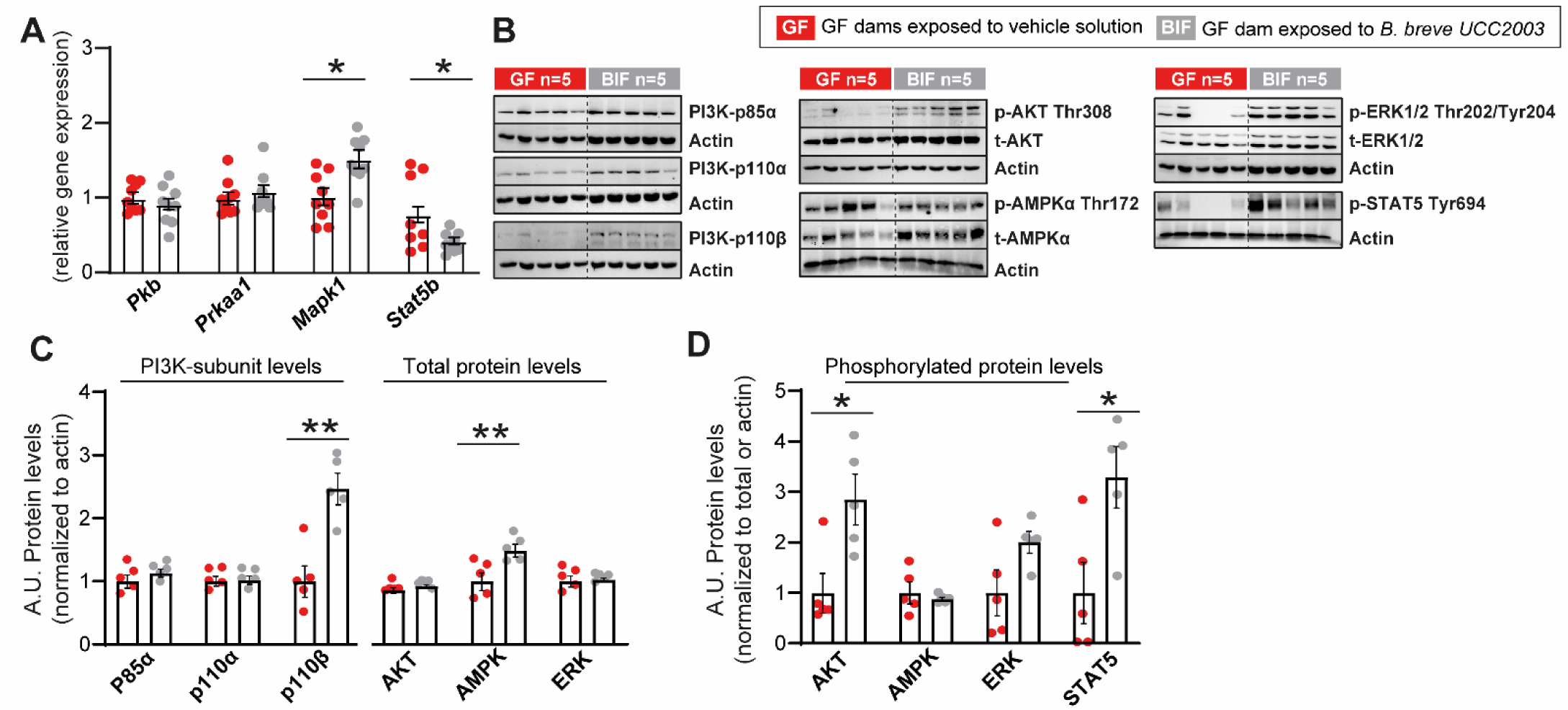
Maternal gut *B. breve* supplementation modifies genes and proteins involved in fetal brain cell metabolism and growth. (A) Relative mRNA levels of genes encoding cell signalling pathways measured in fetal brain samples, expression is relative to two housekeeping genes (*Gapdh* and *Actb*). (B-F) Immunoblots and relative protein expression values of proteins involved in cellular metabolism and growth. Data analysed by ANOVA with the group as fixed effect and means comparisons was made by Fisher test (further details in the methods section). Data are means±SEM with individual datapoints shown. \**P* < 0.05; \*\**P* < 0.01.

### Maternal gut B. breve UCC2003 induces changes in HIF affecting mitochondrial TCA cycle and Wnt-β-catenin signalling in the fetal brain

Molecular pathways like the PI3K-AKT can regulate the stability of hypoxia inducible factors (HIFs) (Joshi et al., 2014). Moreover HIFs regulate brain development and function even in normoxic conditions (Kleszka et al., 2020). We therefore analysed the mRNA and protein levels of HIF-1α and HIF-2α. Whilst we found no difference in the mRNA levels of *Hif1α* or *Hif2α* (Figure 4A), HIF-2α protein abundance was significantly up-regulated in the fetal brain of BIF treated pregnant mice compared to the untreated GF group (no change HIF-1α; Figure 4B). HIF can regulate the tricarboxylic acid (TCA) cycle by repressing mitochondrial function and activating expression of the gene encoding pyruvate dehydrogenase kinase 1 (PDHK1) (Kim et al., 2006). We therefore measured the abundance of PDHK1 in fetal brain lysates and found that total protein content was significantly elevated in the BIF treated group compared to the GF (Figure 4C). Moreover, we quantified mitochondrial ATP production capacity by assessing oxidative phosphorylation (OXPHOS) via analysis of OXPHOS complexes. We were able to detect 4 out of the 5 respiratory chain complexes and found that complex-II was significantly elevated in the fetal brain of BIF treated compared to the untreated GF mice. No changes were observed in the levels of other OXPHOS complexes detected (Figure 4D).

**Figure 4.**
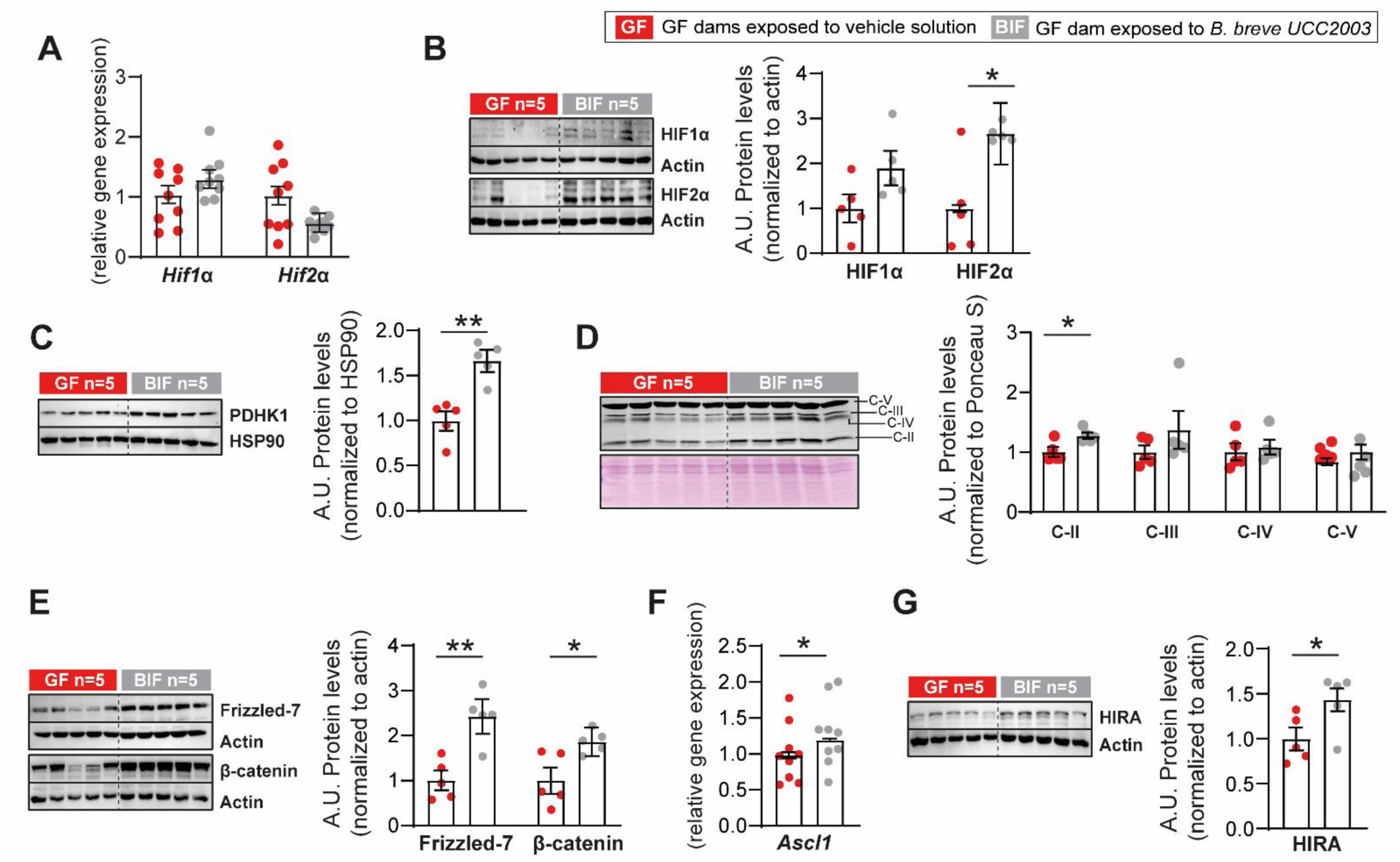
Maternal gut *B. breve* supplementation results in stabilization of HIF2α and changes in mitochondrial function and Wnt-β-catenin signalling in the fetal brain. (A) Relative mRNA levels of *Hif1α* and *Hif2α.* (B-C) Immunoblots and relative protein expression values of HIF1α, HIF2α and PDHK1 normalized to actin or HSP90 levels. (D) Immunoblot and relative protein abundance of mitochondrial complexes (C) normalized to Ponceau Staining. (E) Immunoblots and relative protein abundance of Frizzled-7 and β-catenin normalized to actin levels. (F) Relative mRNA levels of *Ascl1*. (G) Immunoblot and relative protein abundance of HIRA. All gene expression is relative to two housekeeping genes (*Gapdh* and *Actb*). Western blotting performed with 5 fetuses/group from 5 dams per group (different litters). Data analysed by ANOVA with the group as fixed effect and means comparisons was made by Fisher test (further details in the methods section). Data are means±SEM with individual datapoints shown. \**P* < 0.05; \*\**P* < 0.01.

HIF is also known to activate multiple genes and signalling cascades, including the Wnt and β-catenin pathway (Dunwoodie, 2009; Shen et al., 2022). We therefore quantified the levels of proteins in these pathways and observed that the abundance of Frizzled-7, a cognate Wnt receptor (Phesse et al., 2016) and total levels of β-catenin protein were significantly elevated in the fetal brain of BIF treated GF mice (Figure 4E). Moreover, the abundance of *Ascl1*, a pro-neural transcription factor that also induces the Wnt signalling pathway (Tenjin et al., 2019; Woods et al., 2022) was significantly increased in the brain of fetuses from BIF treated GF mice compared to the untreated GF group (Figure 4F). Finally, the abundance of histone cell cycle regulator (HIRA), a histone chaperone that regulates neurogenesis by enhancing β-catenin expression (Li and Jiao, 2017), was enhanced in the fetal brain of GF mice treated with BIF versus untreated GF mice (Figure 4G). Taken together, these results suggest that the presence of *B. breve* in the maternal gut stabilizes HIF2-α, enhances mitochondrial oxidative phosphorylation capacity and promotes Wnt-β-catenin signalling in the fetal brain.

## Discussion

Previous studies have established that the gut microbiome can influence brain function and behaviour (Collins et al., 2012; Sherwin et al., 2019), including a recent murine model study revealing that the maternal gut microbiota can modulate fetal axonogenesis via microbially regulated metabolites (He et al., 2020). These data suggest that targeting the maternal gut microbiota, such as through probiotic supplementation, may lead to defined and specific changes in brain development across the perinatal period. Numerous species and strains of *Bifidobacterium* are commonly found in healthy breast-fed infants, as well as being used as probiotics, where they exert a range of beneficial health effects in the host, including protection against inflammatory insults and promoting barrier function and differentiation of intestinal epithelial cells (Hughes et al., 2017; Kiu et al., 2020b). In terms of brain function, prior work has demonstrated that strains of *Bifidobacterium,* like *B. longum* 1714™, can modify brain activity by improving stress responses and cognitive function in human and animal models (Allen et al., 2016; Savignac et al., 2014; Savignac et al., 2015; Wang et al., 2019). In the current study, we report that oral administration of *B. breve* UCC2003 to pregnant mice can induce changes in the developing fetal brain, namely through modifying fetal brain metabolism, and genes and signalling pathways involved in cell growth and axonogenesis. Our results have potential translational implications, as around 14% of pregnant women use probiotics during pregnancy in European countries like the Netherlands (Rutten et al., 2016). However, the utilization of probiotics in clinical and nutritional contexts, particularly during pregnancy, remains a topic of continual debate. Thus, to comprehensively understand their influence on the developing offspring, it is essential to define the mechanisms supporting their beneficial effects.

In this present investigation, our data suggest that *B. breve* UCC2003 strongly influences fetal brain metabolism. Notably, the fetal brain of the BIF group exhibited a significant reduction in ten specific metabolites. These findings are particularly intriguing, as our previous study did not reveal any changes in the concentrations of these metabolites in the fetal liver, thus highlighting that *Bifidobacterium* supplementation induces distinct responses in different fetal tissues (Lopez-Tello et al., 2022). Among the metabolites, multiple amino acids were reduced, suggesting that these amino acids, including alanine, leucine and valine may have been used by the fetus. Moreover, other amino acids like arginine or carnitine have previously been reported to be impacted by the maternal microbiota, as shown in studies comparing fetal brains of GF versus specific pathogen free mice (Pessa-Morikawa et al., 2022). Leucine, which is transported by the *Slc7a5* (elevated at gene level in the BIF group) and regulated by MTORC1 (Torigoe et al., 2019), has been shown to be regulated by fetal glycaemia, as glucose concentration decreases leucine oxidation independent of insulin (Liechty et al., 1993). Our previous work showed that administration of *B. breve* UCC2003 resulted in upregulation of the *Slc2a1* transporter in the placental labyrinth zone and in the fetal liver. Moreover, fetal glycaemia was improved in the BIF group and compared to the specific-pathogen-free fetus (Lopez-Tello et al., 2022). Although in this study we did not observe changes in the glucose concentrations in the fetal brain, we found that the abundance of key glucose transporters, specifically, *Slc2a1*, *Slc2a3* and *Slc37a1*, were increased in the BIF group. Interestingly, the mRNA levels of *Slc2a3* and *Slc37a1* in the fetal liver were unaltered in the BIF group compared to GF mice (Lopez-Tello et al., 2022), highlighting once again the different metabolic responses of the fetus based on tissue type. Future work should explore the role of fetal glycaemia and its impact on the metabolism of amino acids like leucine in the different organs of the fetus.

Another metabolite that was reduced in the BIF group was 3-hydroxybutyrate. This was in line with the lower levels of valine, as 3-hydroxybutyrate serves as a crucial intermediate in the metabolism of branched-chain amino acids like valine (Xu et al., 2023). Previous studies indicate that 3-hydroxybutyrate, which can be used as an alternative energy source for the brain and other tissues when glucose becomes scarce (Newman and Verdin, 2017), is highly involved in mitochondrial function as it can modify mitochondrial membrane permeability transition (Kim et al., 2015). Moreover, it is known that neurons treated with 3-hydroxybutyrate exhibit an increase in mitochondrial respiration (Kim et al., 2015). We also found two additional metabolites that were reduced in the BIF group, namely carnitine and citrate, which are involved in the tricarboxylic acid (TCA) cycle. Carnitine plays a crucial role in the transport of long-chain fatty acids into the mitochondria (Ferreira and McKenna, 2017), whilst citrate is an intermediate of the TCA cycle involved in nicotinamide adenine dinucleotide metabolism (Icard et al., 2021). Analysis of mitochondrial electron transfer system (ETS) components revealed that the fetal brains of BIF treated mothers had increased levels of the mitochondrial complex-II. Both citrate and 3-hydroxybutyrate are well known inhibitors of succinate dehydrogenase (mitochondrial complex-II) (Hillar et al., 1975), suggesting a potential link between these results. Moreover, we found that the mRNA levels of the transcription factor *Foxm1* and cyclin *Cdk1* were reduced in the BIF group. Prior work has shown that *Foxm1* and *Cdk1* can also control mitochondrial oxygen consumption rates and mitochondrial abundance (Kobiita et al., 2023). Future experiments could be undertaken to understand the potential interplay between these metabolites and the elevated mitochondrial ETS complex II activity, such as using fetal brain explants or cerebral organoids cultured with and without these metabolites, and using high-resolution mitochondrial respirometry analysis. Moreover, aside from *in vitro* studies to improve our understanding of the mechanisms of action, animal studies using additional control experimental groups are needed (Lopez-Tello et al., 2022).

In our study, we did not observe changes in the mRNA levels of well-known genes involved in angiogenesis or apoptosis, such as *Vegf*, *Tp53*, *Casp3* or *Bax*. However, we observed reduced mRNA levels of *Plxna3.* This gene, controlled by the maternal microbiota (He et al., 2020), encodes a class 3 semaphorin receptor that regulates multiple neurodevelopmental processes including axonal growth and guidance (Sakurai et al., 2012; Steele et al., 2022), and neuronal death (Ben-Zvi et al., 2008). Semaphorins have been shown to interact with critical pathways involved in cell proliferation, growth, and apoptosis including the PI3K/AKT pathway (Castro-Rivera et al., 2008). Brains from the BIF group showed activation of the PI3K pathway as evidenced by increased protein levels of PI3K-p110β and elevated phosphorylation levels of AKT. Activation of this signalling pathway has been linked to enhanced dendritic branching, cellular proliferation, neuronal hypertrophy (Wang et al., 2017). Moreover, the PI3K/AKT has been implicated in the regulation of MCTs (Pérez-Escuredo et al., 2016; Zhang et al., 2018), which were found to be elevated in the BIF group. In our study it is unclear if the administration of this bacterium induced structural changes in the fetal brain and if so, whether the PI3K/AKT pathway could be driving such changes. Therefore, further experiments using structural/histological analysis are needed in order to determine changes in cell division, cell death, and axon development.

Prior research in mice has demonstrated the crucial role of the HIF-2α in facilitating proper brain formation, neural network development and the migration of neural stem cells (Kleszka et al., 2020; Leu et al., 2021). Additionally, HIF-2α has been identified as having protective functions for neural stem cells, promoting neurogenesis, and regulating angiogenic and apoptotic processes (Lopez-Barneo et al., 2001). In our study, we observed that *B. breve* UCC2003 administration led to the stabilization of HIF-2α in the fetal brain without inducing changes in HIF-1α protein stabilization. The unaltered state of HIF-1α in this model suggests the absence of intrauterine hypoxic conditions in the BIF group. In this regard, previous research has indicated that HIF-2α, but not HIF-1α, is present in the brains of adult mice under normoxic conditions, and that HIF-1α protein stabilization occurs only under hypoxic conditions (Kleszka et al., 2020). However, other authors suggest that the stability of HIF-2α protein is similar to that of HIF-1α and relies on oxygen-dependent degradation (Loboda et al., 2010). Following this line of research, *in vitro* work using neonatal rat neuronal cultures has demonstrated that hypoxia causes up-regulation of AMPK. This metabolic signalling pathway aside from acting as a metabolic sensor regulating glycolysis, has neuroprotective and pro-apoptotic effects (Muraleedharan and Dasgupta, 2022). In our work we found that total levels, but not phosphorylated AMPK, were increased in the BIF group. Moreover, additional pathways that have been associated with HIFs and found dysregulated in this model, notably Wnt/β-catenin signalling and STAT5. Indeed, the activity of Wnt/β-catenin pathway is closely related with low oxygen levels (Mazumdar et al., 2010) and STAT5 is a target gene of HIF-2α in hematopoietic stem cells (Fatrai et al., 2011). Lastly, we found that PDHK1 was significantly elevated in the BIF group. It is known that HIF can upregulate the expression of PDHK1, which indeed was found elevated in the fetal brain of the BIF group. This adaptive mechanism is used to sustain energy production by inhibiting the conversion of pyruvate to acetyl-CoA, and promoting glycolysis over oxidative phosphorylation (Erdem et al., 2022). Other studies have found that HIF-2α protein is inhibited or knocked down, the expression of *Ascl1* (which was increased in the BIF group) is enhanced; resulting in the induction of sympathetic nervous system differentiation marker genes (Pietras et al., 2009). Taken together, our data suggest that oxygen levels may be altered in fetuses exposed to *B. breve* UCC2003 when compared to GF untreated mice. Although we could not measure oxygen levels at the tissue level in this model to verify whether the fetuses were hypoxic or not, we can also speculate another potential mechanism linked to the HIF-2α protein stabilization that may relate to changes in the transport region of the placenta (Lopez-Tello et al., 2022). In this regard, we found that *B. breve* UCC2003 oral administration resulted in a thinner placental barrier thickness, and this reduction would be expected to aid in the diffusion of oxygen from the mother to the fetus (Woods et al., 2018). Therefore, it is less likely that the fetuses of BIF pregnant mice are hypoxic, but further work is certainly required.

In conclusion, through the use of germ-free mice as a proof of concept, our previous and current studies have underscored the pivotal role of gut microbes, specifically, *B. breve* UCC2003, in the control of metabolic and cellular pathways in the placenta and in the fetal liver and fetal brain (summarized in Figure 5). While the precise mechanisms governing the alterations induced by *B. breve* UCC2003 necessitate further exploration, our study provides clear evidence that maternal oral intake of probiotics can influence fetal organogenesis.

**Figure 5.**
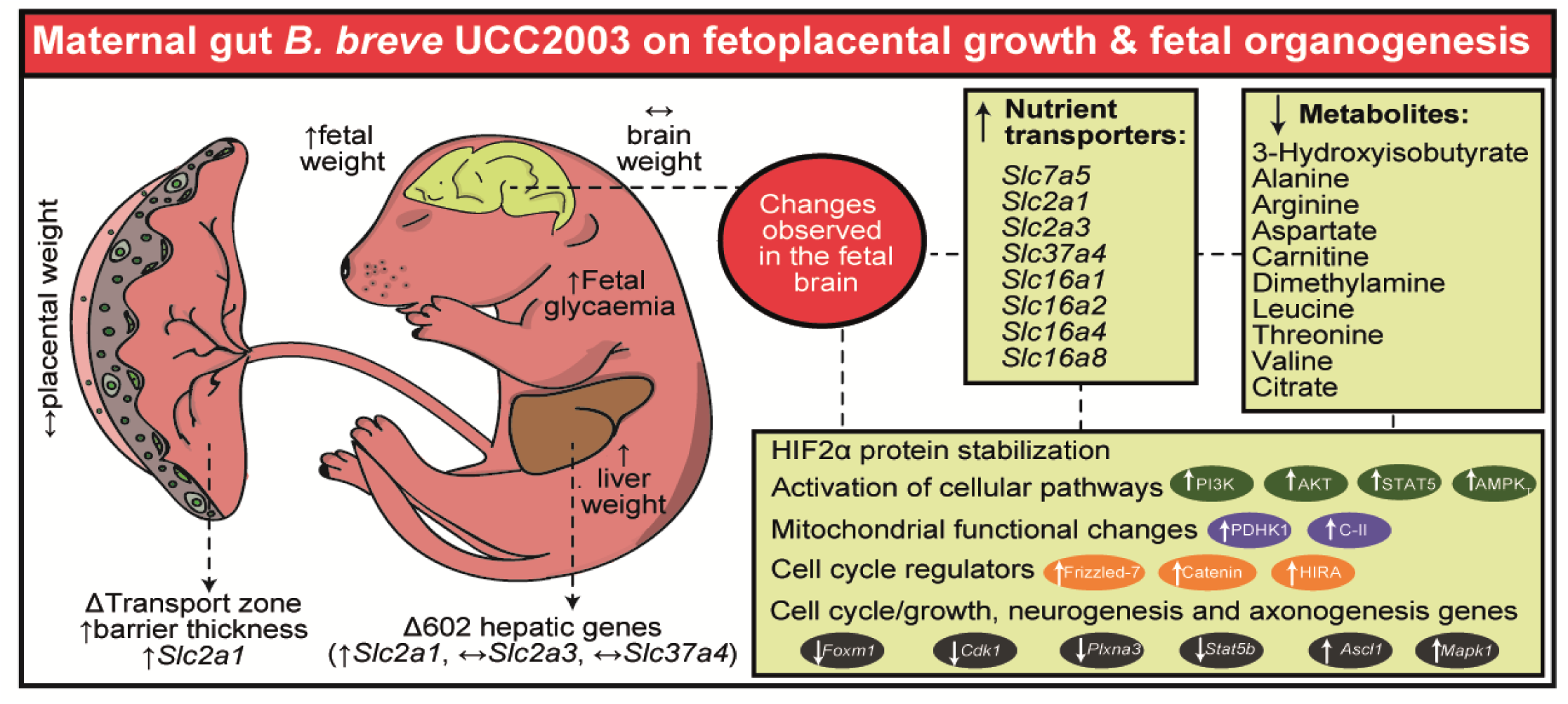
Summary Figure illustrating changes induced by Maternal gut *B. breve* supplementation. The illustration summarizes the most relevant results obtained from our previous study (Lopez-Tello et al., 2022) and the new study comparing GF versus GF treated with *B. breve* during development. C-II refers to the mitochondrial complex-II OXPHOS.

## Limitations of the study

While we have identified numerous changes in gene and protein expression, discerning how these changes impact on the structural organization of the fetal brain following maternal administration of *B. breve* UCC2003 remains ambiguous. Moreover, we used whole brain lysates, resulting in a very heterogenous cellular composition, which limited our ability to study the impact of *B. breve* UCC2003 on cell-type-specific expression in the genes measured. It is noteworthy that our study did not assess the impact of fetal sex on the effects of maternal BIF treatment on fetal development. Additionally, the inclusion of specific-pathogen-free mice in the study, as we did previously (Lopez-Tello et al., 2022), would have provided valuable information to comprehend the direction of the metabolic changes observed in the BIF group. This is important to highlight as the exposure to a microbial challenge in the GF mouse could have also resulted in elevated immune responses. In this regard, it is plausible that the metabolite levels in the BIF group could be within normal ranges for the gestational day and for fetal brain development of the specific-pathogen-free mice (rather than the metabolites being used up more in the BIF treated compared to untreated GF mice). Conversely, untreated GF fetuses might increase the abundance of these metabolites as a mechanism to preserve fetal brain development and maturation in response to growth restriction. For instance, in growth-restricted piglets, there is an increased activity of aromatic amino acid decarboxylase (Bauer et al., 2001). Additionally, the amino acid alanine, which our study found to be altered, is elevated in rats experiencing fetal growth restriction (Chanez et al., 1993). A more sensitive mass spectrometry method may also allow neurotransmitter homologues like GABA to be assessed. This is relevant as there are some data that indicate *Bifidobacterium* can produce these and would be relevant for interpreting our findings (Duranti et al., 2020). Furthermore, the lack of postnatal investigations, including behavioural studies, has constrained our ability to ascertain potential implications of *B. breve* UCC2003 on both short- and long-term neurocognitive outcomes in offspring.

## Materials and Methods

### Bacterial strain, growing conditions and lyophilization

*B. breve* UCC2003 generation/growing conditions were previously described (Lopez-Tello et al., 2022). Briefly, *B. breve* was grown in De Man, Rogosa and Sharpe agar (MRS) under anaerobic conditions overnight. Bacterial cell pellet was resuspended in 10% milk powder and lyophilised in 200ml volume. Lyophilised *B. breve* was reconstituted with 500μl PBS. Concentration of *B. breve* was 10^10^CFU/ml and all batches were tested for contamination upon reconstitution on Luria-Bertani (LB) and Brain-Heart Infusion (BHI) plates under anaerobic and aerobic conditions at 37°C.

### Animal work ethics statement

All animal work was conducted at the University of East Anglia (UK) under the UK Regulation of Animals (Scientific Procedures) Act of 1986 and approved by the UK Home Office and the UEA Ethical Review Committee (project license PDADA1B0C).

### Animal model and experimental design

The animal model employed for this study was previously described (Lopez-Tello et al., 2022) and the experimental design is depicted in Figure S1. Briefly, GF mice were housed under a 12:12 hour light/dark with free access to food and water. GF-C57BL/6J mice were time mated and on gestational day (GD) GD9.5, pregnant GF mice were transferred to individually ventilated cages. Treatment with the probiotic started on GD10 (+2 extra doses on GD12 and GD14) by providing 100µL of reconstituted lyophilised *B. breve* UCC2003 or 100μL vehicle control (PBS, 4 % skimmed milk powder) by oral gavage. The group that received the three doses of *B. breve* UCC2003 received the name of BIF group (n=6 pregnant mice), and the other cohort of mice (treated with vehicle solution) was named as GF group (n=5 pregnant mice). As previously described, our probiotic was formulated in a similar fashion to commercial probiotics preparations. Moreover, the time frame was selected to reflect a potential frame in which women would take probiotics once their pregnancy is confirmed. The colonisation levels of *B. breve* in pregnant mice can be found in our previous publication (Lopez-Tello et al., 2022).

### RNA extraction

Fetal brain RNA was extracted with RNeasy plus Mini Kits (Qiagen) as previously described (Lopez-Tello and Sferruzzi-Perri, 2023). Reverse transcription was performed using the cDNA reverse transcription kit (Applied Biosystems) following manufacturer’s instructions. Samples were analysed with a StepOne real-time PCR machine (ThermoFisher) in duplicates using SYBR Green qPCR master mix (Applied Biosystems, ThermoFisher). Gene expression was normalized to the geometric mean expression of two reference genes, *Actb* and *Gapdh*. Analysis was performed using the 2-ΔΔCt method(Livak and Schmittgen, 2001) and the primer sequences used can be found in Table S2.

### Protein extraction and western blotting assays

Brain protein extraction and Western blotting were performed as previously described (Lopez-Tello and Sferruzzi-Perri, 2023). Briefly, brain samples were lysed with radioimmunoprecipitation assay (RIPA) lysis buffer (R0278-50M, Sigma Aldrich) supplemented with protease inhibitor cocktail mix (11836170001, Roche), 1mM β-glycerophosphate (G-9891, Sigma Aldrich) and 1mM sodium orthovanadate (S65089891, Sigma Aldrich). After protein quantification with BCA protein assay kit (23225, ThermoFisher), samples were mixed with SDS gel loading buffer (L-4390, Sigma Aldrich) and protein denaturalization performed at 90°C for 5 minutes. After electrophoresis, membranes were blocked with 5% fetal bovine serum (A2153-100G, Sigma Aldrich) or semi-skimmed milk (Marvel) and incubated overnight with primary antibodies described in Table S3. The day after, membranes were incubated with secondary antibodies conjugated with horseradish peroxidase (HRP) (1:10,000 NA934 or NA931, Amersham) and exposed to ECL substrate (SuperSignal West Femto, ThermoFisher) for chemioluminiscence detection. Images were taken with the iBright instrument (ThermoFisher) Pixel intensity of protein bands was analysed with ImageJ software.

### Metabolite extraction and nuclear magnetic resonance (NMR) spectroscopy

Fetal brain metabolites were extracted as described elsewhere (Lopez-Tello et al., 2022). Briefly, frozen tissue ∼14 mg was mixed with 200 μL of ice-cold methanol (Fisher Scientific) and 42.5 μL of ultra-pure cold water. Samples were vortexed and tissue was disrupted with a a tissue lyser (Qiagen) and ∼ 15–20 glass beads (Merck) for for 2 × 2 min. Subsequently, 100 μL of ice-cold chloroform (Merck) and 100 μL of ultra-pure cold water were added to the mixture. Samples were incubated on ice for 15 min and then transferred into sterile microcentrifuge tubes and centrifuged for 3 min at 17,000×*g*. The aqueous phase was transferred into new tubes and speed-vacuumed for 30 min at 50 °C and 30 min without heating prior to reconstitution with phosphate buffer solution at 600 μL. Samples were subjected to NMR spectroscopy. The 1H NMR spectra were recorded at 600 MHz on a Bruker AVANCE spectrometer (Bruker BioSpin GmbH, Germany) running Topspin 2.0 software. Fetal brain metabolites were quantified with the software Chenomx® NMR Suite 7.0™.

### Statistical analysis

All statistical analyses and sample sizes are shown in each figure legend. Only samples from viable fetuses were analysed. Statistical analysis was performed with GraphPad Prism software (GraphPad v9, San Diego, CA), SAS/STAT 9.0 (Statistical System Institute Inc. Cary, NC, USA) and Microsoft Excel (v2010). Identification and removal of outliers was performed with the ROUT method (Motulsky and Brown, 2006). For parameters involving only one fetus per litter (Western blot data and metabolomics), a one-way ANOVA was employed with group as a fixed effect and means comparisons were made using the Fisher test (general linear model-GLM model). Conversely, parameters involving more than one fetus per litter (qPCR) were analysed using a one-way ANOVA with the group as fixed effect and each fetus treated as a repeated measure, employing the Fisher test for means comparisons (linear mixed model - MIXED model). In the MIXED model, fetuses coming from the same litter were nested. In both statistical analyses, litter size served as a covariate. The significance threshold for all statistical tests used in this study was set at *p*<0.05. Figures in the manuscript show mean±SEM alongside individual data points (raw data). However, the reported mean±SEM bars have been adjusted for repeated measures and/or litter size. Data were graphed in GraphPad and figure panels were merged with Adobe Illustrator to display corrected mean±SEM and individual dots.

## Data Availability Statement

All relevant data can be found within the article and its supplementary information.

---

**Figure S1.**
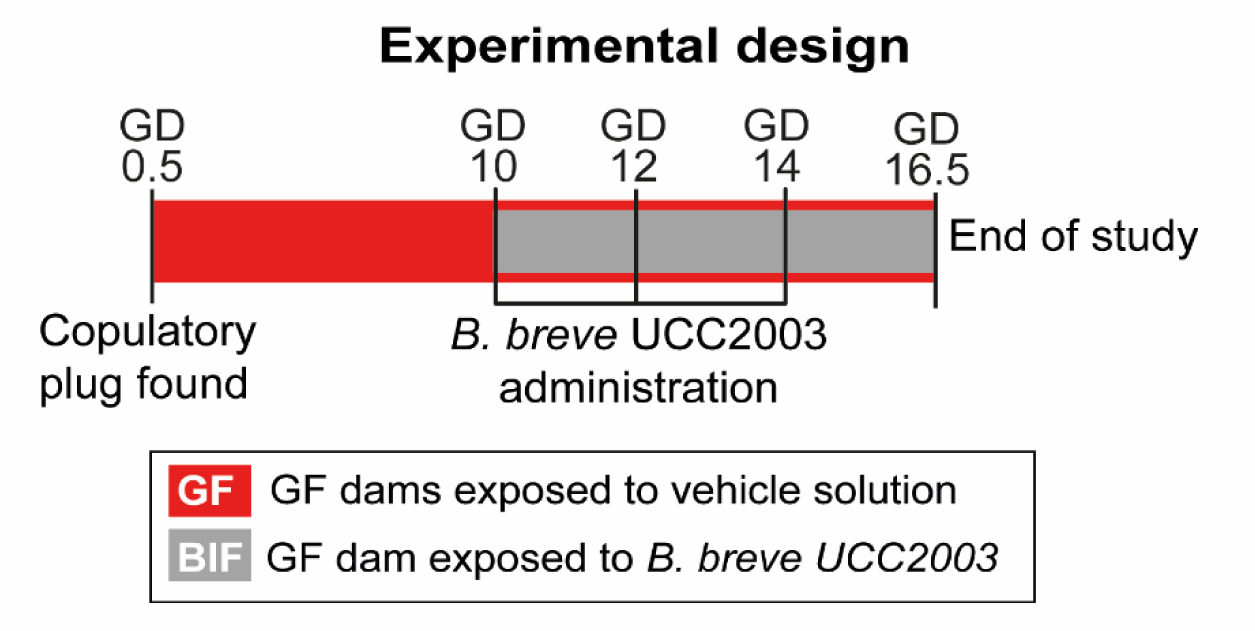
Experimental design. GD: gestational day.

**Table S1. List of metabolites analysed in fetal brains.** Data analysed by one-way ANOVA, with the group as fixed effect and means comparisons made by Fisher test (general linear model-GLM model). Litter size added as a covariate. Data displayed as mean±SEM. Values were considered statistically significant with P<0.05. Additional metabolites can be found in Table 1.

Please see PDF submitted.

**Table S2.**
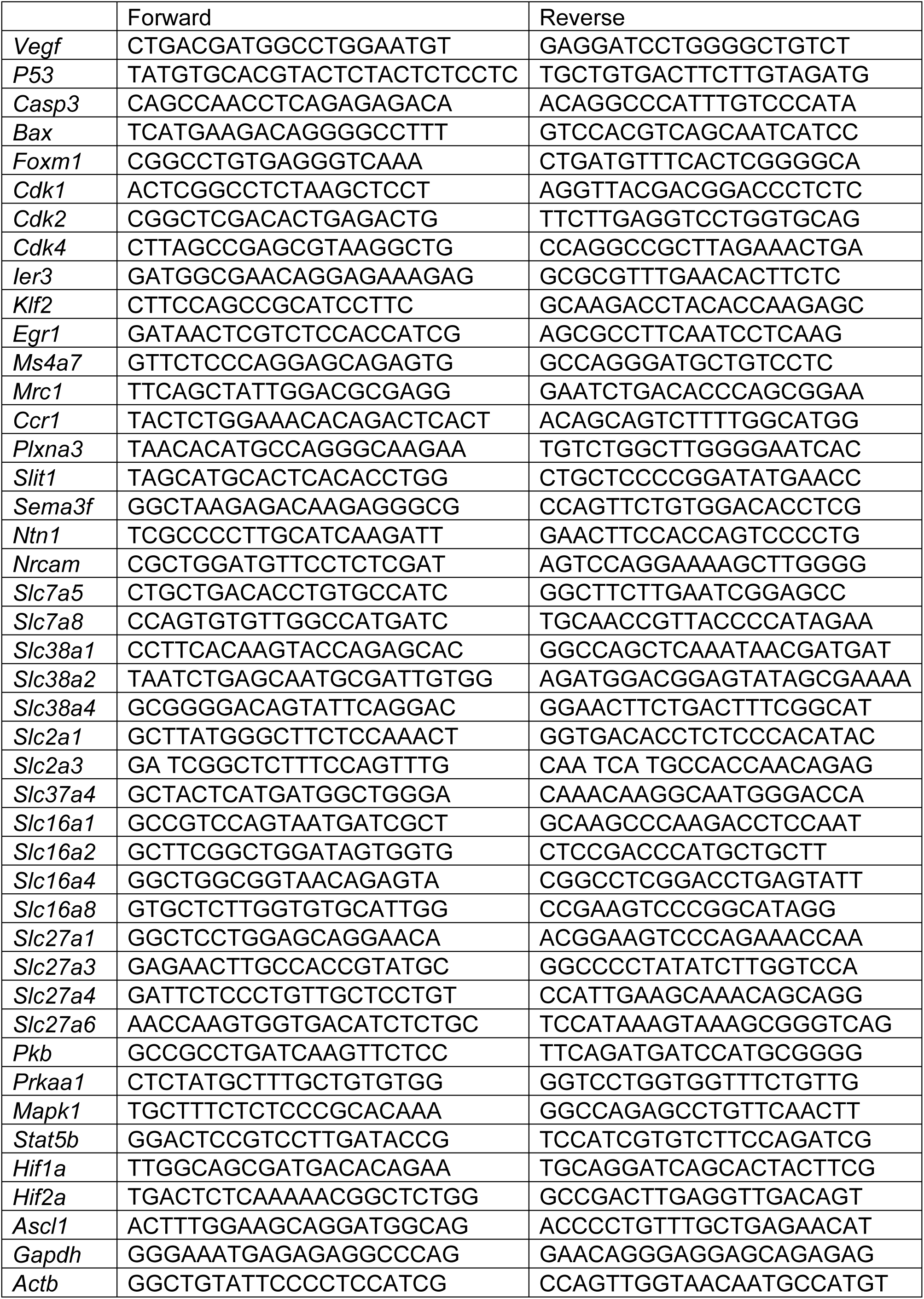
List of primers used for qPCR.

**Table S3.** List of antibodies used for western-blotting.

| Name of antibody | Company | Product number | Dilution |
| --- | --- | --- | --- |
| PI3K-p85 alpha | Millipore | 06-195 | 1:5000 |
| PI3K-p110 alpha | Cell Signaling | 4249 | 1:1000 |
| PI3K-p110 beta | Cell Signaling | 3011 | 1:1000 |
| Total AKT | Cell Signaling | 9272 | 1:1000 |
| Total AMPK | Cell Signaling | 2532 | 1:1000 |
| Total ERK | Cell Signaling | 4695 | 1:1000 |
| Phospho-AKT Thr308 | Cell Signaling | 9275 | 1:1000 |
| Phospho-AMPK Thr172 | Cell Signaling | 2531 | 1:1000 |
| Phospho-ERK Thr202/204 | Cell Signaling | 4370 | 1:1000 |
| Phospho-STAT5 Tyr694 | Cell Signaling | 9351 | 1:1000 |
| HIF1 alpha | Novusbio | NB100-449 | 1:1000 |
| HIF2 alpha | Novusbio | NB100-122 | 1:1000 |
| PDHK1 | Cell Signaling | 3820 | 1:1000 |
| HSP90 | Cell Signaling | 4877 | 1:1000 |
| Frizzled-7 | Abcam | Ab64636 | 1:1000 |
| HIRA | Cell Signaling | 12463 | 1:1000 |
| Actin | Cell Signaling | 3700 | 1:5000 |
| Beta catenin | Bd Biosciences | 610153 | 1:1000 |
| OxPhos cocktail | Thermofisher | 45-8099 | 1:200 |

